# Cell wall proteomics in live *Mycobacterium tuberculosis* uncovers exposure of ESX substrates to the periplasm

**DOI:** 10.1101/2023.03.29.534792

**Authors:** Neetika Jaisinghani, Mary L. Previti, Joshua Andrade, Manor Askenazi, Beatrix Ueberheide, Jessica C. Seeliger

## Abstract

The cell wall of mycobacteria plays a key role in interactions with the environment and its ability to act as a selective filter is crucial to bacterial survival. Proteins in the cell wall enable this function by mediating the import and export of diverse metabolites from ions to lipids to proteins. Accurately identifying cell wall proteins is an important step in assigning function, especially as many mycobacterial proteins lack functionally characterized homologues. Current methods for protein localization have inherent limitations that reduce accuracy. Here we showed that protein tagging by the engineered peroxidase APEX2 within live *Mycobacterium tuberculosis* enabled the accurate identification of the cytosolic and cell wall proteomes. Our data indicate that substrates of the virulence-associated Type VII ESX secretion system are exposed to the *Mtb* periplasm, providing insight into the currently unknown mechanism by which these proteins cross the mycobacterial cell envelope.

## INTRODUCTION

The cell wall of bacteria provides the first line of defense against environmental insults, whether ions, toxins, reactive species, exogenous enzymes, or small molecule inhibitors. Importantly, bacteria must strike a balance between putting up a wall and permitting essential processes—the uptake and efflux of these same classes of molecules—ions, metabolites, proteins—that allow them to survive and grow. The cell wall is thus both barrier and gatekeeper. Barrier function is provided by diverse structures including polymers and noncovalent assemblies of sugars, peptides, and lipids, but gatekeeping is performed largely by proteins that shuttle, span, and otherwise serve as conduits through the cell wall. The cell wall of pathogenic bacteria is also the site of contact with the host. Here, the cell wall plays a key role by presenting specific factors such as lipids and proteins that influence the interaction between pathogen and host and direct infection outcomes. Identifying cell wall proteins that function in essential transport processes, such exporting virulence factors and importing nutrients, is thus a critical step in understanding bacterial pathogenicity and physiology and identifying strategies to combat bacterial infections therapeutically and preventatively.

The problem of transport through the cell envelope is especially challenging in bacteria with two membranes: Cargo must pass through multiple hydrophobic and hydrophilic layers that pose both physical and energetic barriers. Numerous elegant genetic and biochemical studies have uncovered the mechanism of protein, lipoprotein, and lipid transport to and from the outer membrane of gram-negative bacteria^1–4^. However, the pathways that move molecules through the cell wall of mycobacteria, including pathogens like *Mycobacterium tuberculosis*, remain a mystery. Mycobacteria have an odd relationship to their bacterial brethren: Although phylogenetically gram-positive, the structure of their cell envelope has more in common with diderm gram-negatives. In addition to a phospholipid plasma membrane and peptidoglycan layer, mycobacteria have an arabinogalactan polysaccharide layer to which very long-chain fatty acids known as mycolic acids are covalently attached^5^. This mycolate layer is thought to form the inner leaflet of an outer membrane, which is more commonly known as the mycomembrane due to its unique structure and lipid composition^6^. Given the distinct structure of the mycobacterial cell envelope, it comes as no surprise that Gram-negative transport proteins, such as those from the Lol lipoprotein and Lpt lipopolysaccharide systems^2, 7^, have no homologues in mycobacteria.

Irrespective of their homology (or lack thereof), transport proteins must share one universal property: their location in the cell wall, which is a clear hallmark of their function. In general, subcellular localization is an important consideration in predicting protein function, especially when there is no homology to proteins of known function. Determining protein subcellular localization experimentally in bacteria is a particular challenge due to their small size. Mycobacteria require yet more specialized protocols because their cell envelope responds differently to lysis and extraction methods. Physical separation methods require the differential sedimentation of fragments generated by cell lysis^8–11^. Selective extraction methods separate proteins by physiochemical properties based on their interactions with detergents such as Genapol and Triton X-114^12, 13^. Genetic methods use the subcellular compartment-specific activity of fusions to proteins such as GFP, whose fluorophore matures only in the cytoplasm^14^, and beta-lactamase^15^ or alkaline phosphatase^16^, which are active only following export beyond the plasma membrane, and assume that protein localization is not affected in the fusion constructs. Photocrosslinking lipids enable the identification of proteins that interact with these membrane components^17^. Each of these methods have helped localize specific proteins or identify subcellular proteomes, but have considerations that limit their accuracy. For example, differential sedimentation is often used due to its simplicity, but especially with highly sensitive mass spectrometry, detection of abundant cytoplasmic proteins in all fractions is common. In addition, soluble proteins from the periplasm do not necessarily sediment with cell wall fractions and can be misassigned to the cytoplasm. Predicting protein export with empirical algorithms also has limitations: Not all exported proteins have a canonical N-terminal signal peptide. Thus, there has been a need for methods that can identify cell wall proteins more accurately, towards understanding the mechanisms of cell wall processes.

We have previously reported the application of the engineered peroxidase APEX2 to subcellular compartment-specific protein labeling within live mycobacteria^18^. APEX2 is one of several enzyme-mediated protein tagging methods known as proximity labeling^19^, but is particularly well suited to the bacterial cell wall because unlike variants of biotin ligase (*e.g*., BioID^20^, TurboID^21^ and others), it does not require ATP, which is absent from the bacterial periplasm^22^. We showed in the fast-growing species *Mycobacterium smegmatis* that APEX2 labeling can localize endogenously or heterologously expressed proteins to the cytoplasm or cell wall with high accuracy and specificity^18^. Here, we applied this technology to the human pathogen *Mycobacterium tuberculosis*. We first validated the accuracy of localization for examples of known cell wall proteins. We then identified the cytoplasmic and cell wall proteomes tagged by APEX2. Extensive comparison to cell wall proteomes identified by other methods confirmed the higher accuracy of our approach. Most importantly, we detected substrates of Type VII ESX protein secretion system in the cell wall, providing the first experimental evidence that these proteins are exposed to the periplasmic environment, with implications for the model of ESX-mediated protein export through the cell envelope.

## RESULTS

*Optimization and validation of subcellular compartment-specific labeling by APEX2 in* Mycobacterium tuberculosis. We have previously reported optimization of APEX2 for compartment-specific labeling in both *M. smegmatis* and *Mtb*^23^. Briefly, we found that in *Mtb* codon optimization of the APEX2 gene was necessary for export of APEX2 into the cell wall via fusion to the N-terminal signal peptide from the *Mtb* gene *mpt63* (hereafter referred to as Sec-APEX2): Only codon-optimized Sec-APEX2 was enzymatically active in whole cells with the colorimetric substrate guaiacol and yielded a distinct pattern of labeling by the substrate biotin-phenol compared to APEX2 expressed in the cytoplasm (Cyt-APEX2) (**Figure 1A; Figure S1A**). To confirm compartment-specific labeling, lysates were enriched for biotinylated proteins and probed with native antibodies to previously validated representative cell wall proteins: the group of mycolic acid acyltransferases known as the antigen 85 complex (Ag85 or Ag85ABC; also called FbpABC) and the lipid transfer-associated lipoprotein LprG (**Figure 2A**). As predicted, both Ag85 and LprG were enriched from *Mtb* expressing Sec-APEX2, but not Cyt-APEX2. Notably, using subcellular fractionation by differential centrifugation, the Ag85 complex partitions into the soluble fraction^24^, which is often interpreted as the cytoplasm. In contrast, APEX2 labeling clearly distinguishes Ag85 as located in the cell wall.

**Figure 1.**
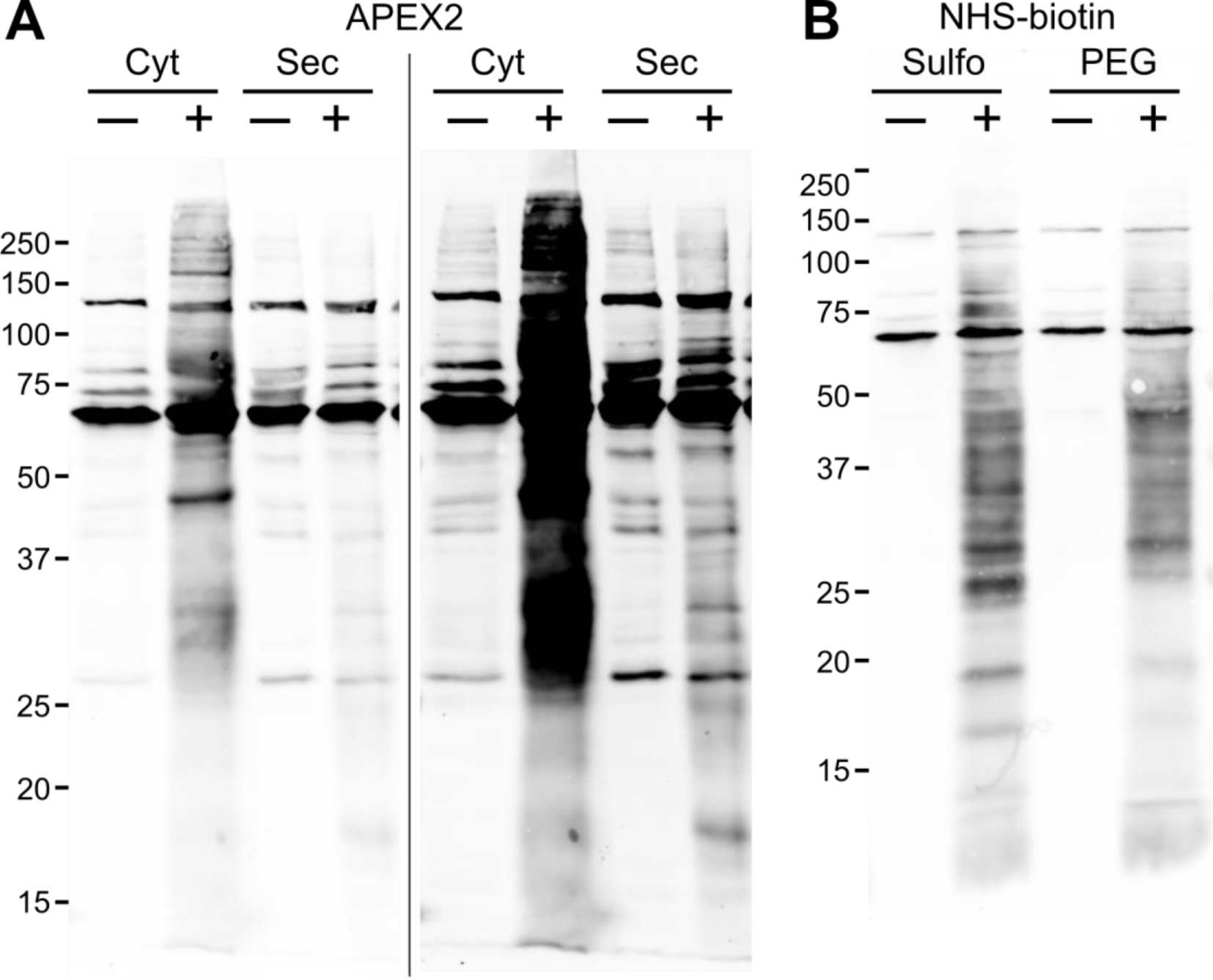
Biotinylation of proteins in Mtb by Cyt-APEX2, Sec-APEX2 or N-hydroxysuccinimide reagents. (A) *Mtb* encoding inducible Cyt-APEX2 or Sec-APEX2 was cultured without (─) or with (+) theophylline and subjected to the labeling protocol with biotin-phenol. Image on the right is the same blot contrasted to highlight Sec-APEX2-dependent biotinylation. (B) *Mtb* whole cells were treated with sulfo-NHS-biotin (Sulfo) or NHS-PEG-biotin (PEG). For all blots, biotinylation in crude lysates were detected with streptavidin. Immunoblots are (A) representative of >3 independent experiments (see Figure S2 for additional replicates and (B) from one experiment for sulfo-NHS-biotin and representative of 3 independent experiments for NHS-PEG-biotin. The fold increase in biotinylation upon induction of APEX2 or treatment with NHS (“Ratio”) was calculated by taking the ratio of the + / − theophylline output intensities after normalizing to the corresponding inputs.

**Figure 2.**
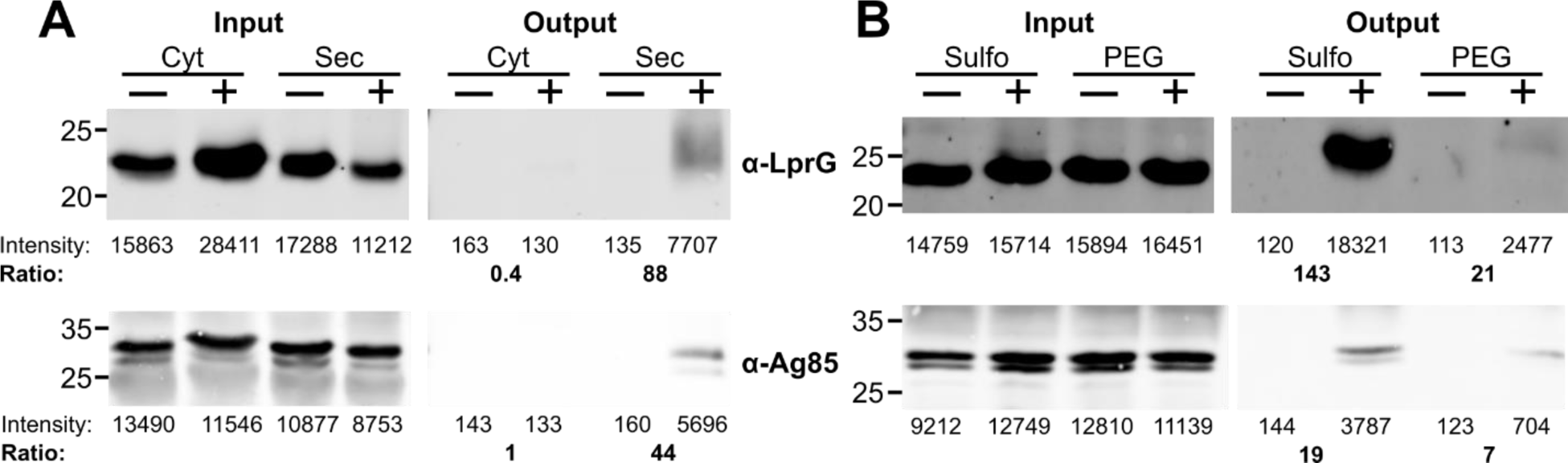
Validation of compartment-specific labeling by Cyt-APEX2, Sec-APEX2 or N-hydroxysuccinimide reagents. Biotinylated proteins were enriched from crude lysates obtained from Mtb treated as follows: (A) Mtb encoding inducible Cyt-APEX2 or Sec-APEX2 were cultured without (─) or with (+) theophylline and subjected to the labeling protocol with biotin-phenol. (B) *Mtb* whole cells were treated with sulfo-NHS-biotin (Sulfo) or NHS-PEG-biotin (PEG). Immunoblots are (A) representative of 2 independent experiments (see Figure S2 for additional replicate) and (B) from one experiment for sulfo-NHS-biotin and representative of 2 independent experiments for NHS-PEG-biotin.

*Cyt-APEX2 and Sec-APEX2 tag distinct subsets of the* Mtb *proteome*. Having confirmed compartment-specific labeling by APEX2, we proceeded to enrich and identify APEX2-dependent biotinylated proteins by label-free quantitative proteomics. We note that *Mtb* natively biotinylates some proteins (**Figure 1**) and acknowledge that unless additional biotinylation by APEX2 significantly increases their enrichment efficiency, these endogenously biotinylated proteins will not be detected in the protocol used here. Although we previously reported alternative APEX2 substrates that do not use biotin and thus avoid this issue, we chose to proceed with biotin-phenol due to existing robust pipelines for affinity purification and downstream proteomics.

We identified proteins as labeled by APEX2 if they were enriched in all 3 experimental replicates upon expression of Cyt-APEX2 (486 proteins) or Sec-APEX2 (254 proteins) compared to uninduced controls (**Table S2**; see Methods Details). For simplicity, we hereafter refer to these as the Cyt and Sec proteomes, respectively. We detected far fewer than the number of predicted proteins in *Mtb* (4,023) or in previously reported total and cell wall proteomes (as many as 3,894 and 1,729, respectively^9, 25^). However, this may be consistent with the selective chemistry of APEX2 labeling, which tags only tyrosines accessible to chemical modification. As expected, the Sec proteome was enriched compared to the Cyt proteome for proteins containing predicted transmembrane helices, signal peptides for the Sec and Tat secretion systems, and/or a conserved lipoprotein motif (lipobox) (**Figure 3; Table S2**).

**Figure 3.**
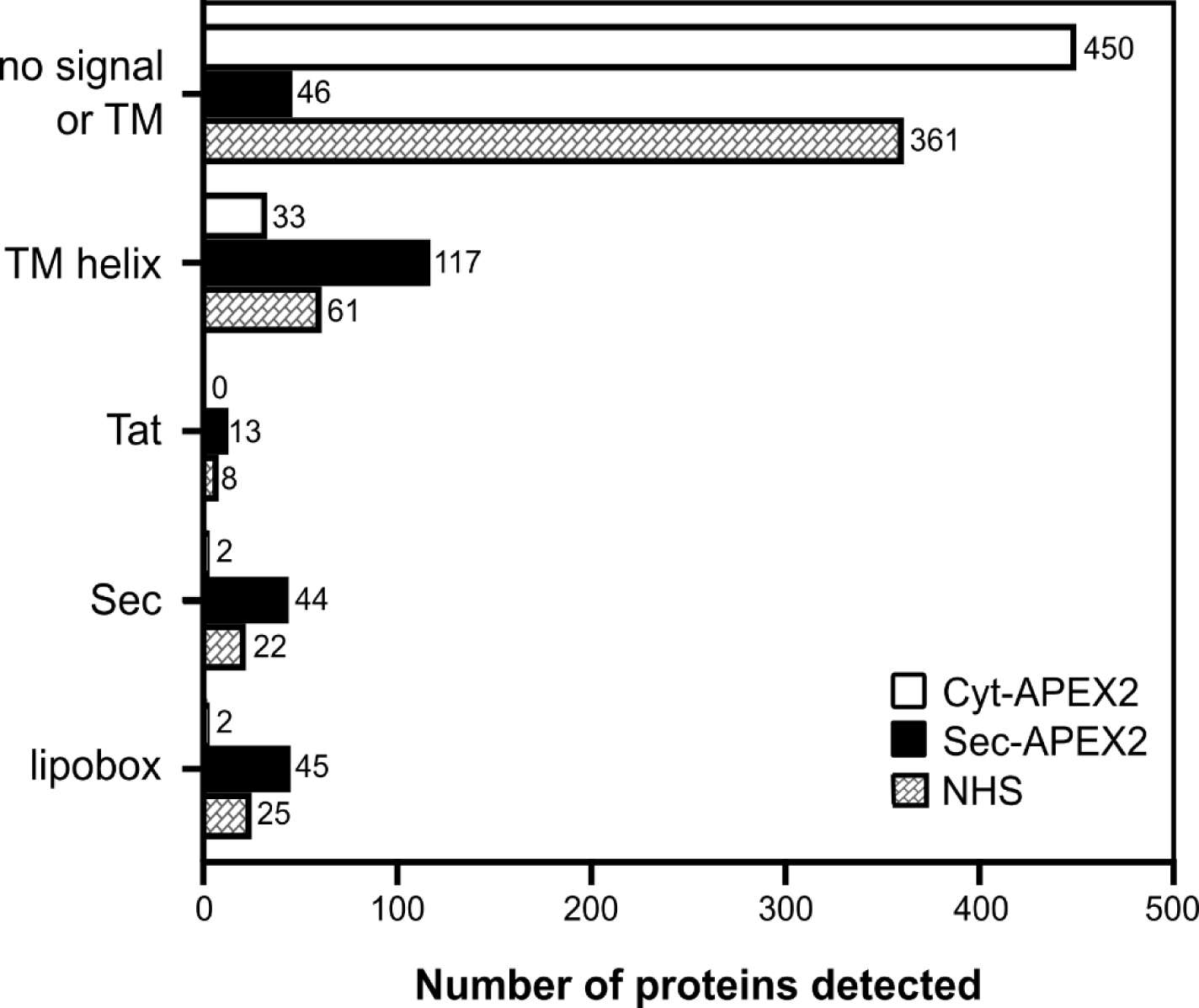
Analysis of Cyt, Sec, and NHS proteomes for predicted signal peptides and transmembrane helices. Protein sequences from the Cyt (486), Sec (254), and NHS (416) proteomes were analyzed using TMHMM and SignalP to detect predicted transmembrane helices and Sec, Tat, and lipobox sequences. Data labels indicate the number of proteins in each category.

*N-succinimidyl ester derivatives of biotin do not have high specificity for* Mtb *surface proteins*. In early experiments we sought to label the cell surface proteome by tagging surface lysines with biotin linked to N-hydroxysuccinimide (NHS). We chose two reagents reported not to cross membranes due to their charge and/or hydrophilicity (sulfo-NHS-biotin, NHS-PEG-biotin). Although both reagents yielded patterns of labeling distinct from either APEX2 or Sec-APEX2 (**Figure 1B, Figure S1B**), the cell wall proteins LprG and Ag85 were labeled by both NHS reagents and with apparently higher efficiency by sulfo-NHS-biotin than NHS-PEG-biotin (**Figure 2B**).

We went on to identify proteins modified by NHS-PEG-biotin vs. vehicle-treated control using the same avidin enrichment and analysis protocol as above. This NHS-tagged proteome (416 proteins; **Table S2**) contained many proteins not identified in either the Cyt or Sec proteomes (114 unique hits). This is not surprising, as the number and accessibility of lysines targeted by NHS vs. tyrosines targeted by APEX2 is likely to differ for each protein across the proteome. However, compared to the Sec proteome, the NHS proteome contained a far higher number of proteins lacking any predicted export signal (361 vs. 46; **Figure 3**) and overlapped exclusively with the Cyt proteome (196 proteins; 47%) far more than with the Sec proteome (77 proteins; 19%) (**Table S2**). Finally, many proteins in the NHS proteome that lacked a predicted export signal or transmembrane helix had predicted or confirmed cytosolic functions, including in the categories of amino acid metabolism (20 proteins) and nucleic acid metabolism (including replication, transcription, translation and DNA repair) (67 proteins) (**Table S3**). Overall, our results show that in mycobacteria, sulfo-NHS-biotin and NHS-PEG-biotin do not label exclusively surface proteins, but rather penetrate the cell envelope to a significant degree on the timescale of the labeling reaction. While altering the conditions of the labeling reaction (incubation time, temperature) may enable more specific labeling, we chose to focus on APEX2 rather than optimize NHS labeling further.

*The Sec-APEX2-tagged proteome shows high selectivity for cell wall proteins*. While our initial analysis (**Figures 2A, 3**) suggested that Sec-APEX2 acts within the cell wall, for rigor we also asked, does Sec-APEX2 tag cytoplasmic proteins? Of the 254 proteins in the Sec proteome, 46 lacked a predicted signal sequence or transmembrane helix (**Figure 3, Table S2**). Of these, 16 proteins are conserved hypothetical proteins with no functional annotation and we did not include them further in our analysis. The remaining 30 are annotated in the functional categories of cell wall and cell processes; intermediary metabolism and respiration; information pathways, regulatory proteins, or lipid metabolism (**Table S4**). While several such as PonA2 are known or predicted to be exported, we cannot eliminate the possibility that the rest are bona fide cytoplasmic proteins. For example, labeling by Sec-APEX2 may result from limited diffusion of phenoxyl radicals outside of the targeted compartment^26, 27^. Indeed, three of these proteins (IspH2, FhaA, PapA1) may be membrane proximal based on their functions or interactions with membrane proteins^28, 29^ and thus more likely to react with Sec-APEX2-generated radicals that cross the membrane (**Table S4**). The other 19 soluble proteins include predicted ribosomal factors and DNA-interacting proteins (helicase, transcription factors, etc.), or enzymes with predicted functions in processing cytoplasmic metabolites. Based on our other strong evidence for compartment-specific labeling, we speculate that these soluble proteins have alternate or moonlighting functions in the cell wall. Regardless, their detection in the Sec proteome will inform their further functional characterization. Overall, we concluded that the Sec proteome reports accurately and specifically on proteins that occupy the cell wall.

We then compared the Cyt and Sec proteomes to those obtained in studies that sought to identify the cell wall or exported proteome in *Mtb*. We focused first on proteomes identified from fragments containing the cell envelope (plasma membrane and cell wall) that are obtained by sedimentation because this method is broadly used to localize both proteomes and individual target proteins^9, 30–32^ (**Table S5**). (Cell wall or membrane-associated proteomes obtained by detergent extraction have also been reported^12, 13^, but for consistency in comparing methods, were not included in our analyses.) As previously noted, abundant cytoplasmic proteins are commonly detected in cell envelope fractions, including proteins involved in amino acid metabolism, nucleic acid metabolism (including replication), transcription, and translation. While many of these proteins were detected in the Cyt proteome, they were notably absent from the Sec proteome (**Table S5**). This analysis confirmed that Sec-APEX2 labeling yielded a higher degree of specificity for the cell wall than the isolation of cell envelope fragments.

APEX2 is not the only genetically encoded strategy for discriminating cell wall proteins in the live mycobacteria; the Braunstein lab has pioneered the use of cell wall-active enzyme insertion fusions on a genome-wide scale to analyze the identity and topology of exported proteins. Their Exported In vivo Technology (EXIT) analysis in mice revealed 593 proteins exported by *Mtb* during infection^33^. As expected, the EXIT and Sec proteomes have significant overlap: Nearly 70% of the Sec proteome (176 of 254) was also identified by EXIT (**Table S6**). Interestingly, 11 proteins detected exclusively in our Cyt proteome were also identified by EXIT, but nearly all have predicted transmembrane helices. This suggests that their identification by cytoplasmic APEX2 labeling is due to the labeling of tyrosines in cytoplasmic domains.

Above we focused on results obtained by methods that localize proteins by their physical association with cell envelope fragments or by the activity of a fusion to a cell wall-active enzyme. Recently, Kavunja et al. identified putative cell wall proteins in *M. smegmatis* by their crosslinking in the cell wall to a photoreactive analogue of the mycomembrane lipid trehalose dimycolate^17^. This study detected 7 orthologues of proteins found also in our Sec proteome: FhaA (Rv0020c), FbpC/Ag85C (Rv0129c), MmpL3 (Rv0206c), MmpL5 (Rv0676c), MarP (Rv3671c), Rv1410c, and Rv0227c. All are established divisome (FhaA), cell wall (FbpC/Ag85C) or integral plasma membrane proteins (MmpL3, MmpL5, MarP, Rv1410c, Rv0027c). In addition, Kavunja et al. detected 18 orthologues that we identified exclusively in the Cyt proteome (**Table S7**).

The detection of FhaA by Kavunja et al. and in the Sec proteome is striking, as noted above its expected cytoplasmic localization has been underscored by its in vitro binding to the juxtamembrane domain of the integral membrane receptor-like kinase PknB^34^ and its co-crystallization with the phosphorylated pseudokinase domain of the peptidoglycan biosynthesis integral membrane protein MviN^35^. As noted above, FhaA is also associated with the divisome and interacts in a 2-hybrid assay with another peptidoglycan biosynthesis protein, PbpA^36^. This led us to consider whether the barrier function of membranes may be compromised at sites of new cell wall synthesis. If this were true, then we would predict that both the Cyt and Sec proteomes would contain multiple proteins that are part of the divisome or elongasome complexes. However, we found that our detection of 12 known divisome/elongasome proteins was largely consistent with expected localization (**Table S8**): one cytoplasmic protein was found exclusively in the Cyt proteome (Wag31, or DivIVA) and 8 known cell wall or integral membrane proteins were found exclusively in the Sec proteome (RipA, PonA1, Pbpa, PbpB/FtsI, PonA2, FtsX, FtsQ, LdtB)^37–41^. The exceptions were FhaA and the predicted cell wall protein LamA^38^, which were found in both Cyt and Sec proteomes, and the septal protein SepIVA, which was found only in the Sec proteome. The cross-compartment detection of these 3 proteins is intriguing and inspires speculation that they have special roles in cell growth and division processes that involve mixing between compartments (*e.g*., membrane fusion).

Identification of APEX2-labeled proteins can also be combined with structural information to analyze membrane protein topology. Our experimental protocol did not enable identification of the biotinylated peptides and so does not provide site-specific information, but topology can be implied for proteins that have accessible tyrosines on only one side of the membrane. This is likely to be the case, for example, for single-pass membrane proteins with one predicted transmembrane helix. Indeed, the peptidoglycan enzymes PbpA, PbpB, PonA1, and DacB are all detected exclusively in the Sec proteome, consistent with their enzymatic domains facing the periplasm and as would be assumed based on their functions (**Figure S2**). On the other hand, the protease PepD (also known as HtrA2; Rv0982) is detected exclusively in the Cyt proteome (**Figure S2**). Our data also establish topology for several conserved hypothetical proteins of unknown function (**Figure S2**).

*Substrates of the Type VII ESX secretion system are exposed to the periplasm*. The Type VII ESX secretion systems are unique protein export systems found within mycobacteria and Firmicutes^42^. Detailed structures are available for the plasma membrane assembly known as the core complex^43–46^ and for various oligomers thought to represent cytoplasmic complexes of chaperones and secretion substrates^47–50^. However, there is still no evidence for an ESX apparatus that spans the cell wall and how proteins are exported across the periplasm and through the mycomembrane remains unknown. The APEX2-labeled proteomes were consistent with the expected topology of the ESX core complex: Core proteins were detected exclusively in the Cyt or Sec proteomes depending on whether their soluble domains were expected to be on the cytoplasmic (EccC2, EccE1) or periplasmic side (MycP1, P2, P3, P5; EccB1, B2, B3) of the plasma membrane, respectively (**Table S2**). The Cyt proteome also included other Esx components predicted to be in the cytoplasm, including the putative ESX-1 chaperones EspL and EspH, which are required for the stability and secretion of EspE and EspF^51, 52^. Finally, the ESX-1 substrates EsxB, EspB, and EspK were also in the Cyt proteome, consistent with their previous detection in total cellular lysates^53–55^, as well as the substrates EsxL/EsxN (due to redundancy in tryptic peptides, these two proteins could not be uniquely assigned).

Given the demonstrated consistency and specificity of APEX2 labeling, detection of the ESX substrates EsxB and EsxG in the Sec proteome was striking. EsxB (also known as CFP-10) and EsxG are homologues of each other and substrates of the ESX-1 and ESX-3 secretion systems, respectively. Based on co-crystal structures with their complementary ESX substrates EsxA (ESAT-6) and EsxH, tyrosines on EsxB (Y83) and EsxG (Y83, Y94) could be surface accessible^49, 56^. Our results provide direct evidence that tyrosines in EsxB and EsxG are labeled by phenoxyl radicals generated by APEX2 and is thus exposed to the periplasmic environment.

*Proteins in* Mtb *culture filtrates are a mixture of cytosolic and cell wall proteins*. As presented above, multiple lines of evidence indicate that the Sec proteome accurately reports on proteins that are resident in the cell wall, including proteins that may be in transit to the cell surface or extracellular milieu. Therefore, we also analyzed the overlap between the Sec proteome and proteomes detected from culture filtrates^57–60^. Proteins released into the medium from *Mtb* cultures (culture filtrates) are often classified as secreted, although released proteins may also originate from lysed cells or surface shedding due to detergent typically included in the growth medium. We asked whether localization of proteins by APEX2 tagging could validate the culture filtrate proteome as the secretome. We compared the Cyt and Sec proteomes to 4 culture filtrate proteomes (**Table S9**). Of the proteins detected in ≥3 studies, nearly half (191 of 418) were detected by APEX2 labeling. Of those, 99 (52%) were detected exclusively in the Cyt proteome and 80 (42%) in the Sec proteome (12 proteins, or 6%, were detected in both proteomes). Importantly, culture filtrate proteins detected only in the Cyt proteome have functions in cytoplasmic processes and are predicted to be released only upon cell lysis. These include 10 involved in amino acid metabolism and 19 with functions in replication, transcription, or translation, including the RNA polymerase subunit RpoA. Of the 227 culture filtrate proteins not detected by APEX2 labeling, 204 (∼90%) had no predicted signal peptide or transmembrane helices. This analysis suggests that culture filtrate proteomes contain significant numbers of cytoplasmic proteins that may be stable and/or abundant species detected even when only a low level of cell lysis is expected.

## DISCUSSION

Our results established APEX2-mediated protein tagging as a highly accurate method for localizing proteins to the cytoplasm and cell wall of mycobacteria. The key limitation is that target proteins must have a tyrosine with sufficient exposure to react with the phenoxyl radical generated by APEX2, but our detection of EsxB, for example, shows that the presence of even a single accessible tyrosine can be sufficient. Another potential limitation is that since labeling is performed in live cells, proteins in macromolecular complexes may be excluded. For example, we did not detect any of the 12 proteins from the multi-protein fatty acid synthase II (FAS-II) complex (Rv1482c, Rv1483, Rv1484, Rv1485, Rv2243, Rv2244, Rv2245, Rv2246, Rv2247, Rv0635, Rv0636, Rv0637), even though they have been detected in total *Mtb* proteomes^25^ and all have tyrosines. We also note that phosphorylation of tyrosine residues is unusually prevalent in *Mtb*^61^ and could also limit tyrosines available for tagging. Finally, as with all methods, detection is also limited by protein expression; more comprehensive subcellular proteomes may be achieved with APEX2 labeling performed in additional strains and under additional culture conditions in which individual protein levels are altered.

In curating the APEX2-tagged proteomes, the identification of three cell growth/division proteins (SepIVA, LamA, FhaA) in the cell wall raises intriguing questions. First, these are presumed cytoplasmic proteins based on the absence of a predicted N-terminal signal peptide. If they are indeed in the cytoplasm, does APEX2-mediated tagging occur across subcellular compartments because the cytoplasm and periplasm intermix at some point, either within the cell or during the cell cycle, e.g., at septa during cell division? Do these proteins concentrate at such points because they have roles in mediating processes that require membrane fusion or mixing? Second, all three are phosphorylated^62^ and/or arginine methylated^63^ at multiple sites, presumably in the cytoplasm. However, a recent comprehensive study of the *Mtb* O-phosphoproteome uncovered unique phosphosites on many predicted and validated cell wall proteins, suggesting that phosphorylation is not limited to cytoplasmic proteins^61^.

The mechanism of ESX secretion through the cell wall is unknown. Substrates EsxB and EsxG are exported by the ESX-1 and ESX-3 secretion systems, respectively. Our detection of substrates from two of the five ESX systems found in mycobacteria suggests that exposure to the periplasm is a general feature for these homologues. EsxB and EsxG form stable complexes with EsxA and EsxH, respectively. Because EsxA and EsxH are not detected in our cell wall proteome, it is tempting to speculate that their accessible tyrosine(s) are engaged in specific interactions that mask them from phenoxyl radicals. On the other hand, while we used a stringent and accepted SAINT score cutoff of 0.90 to identify hits that were enriched in APEX2-expressing vs. non-expressing *Mtb*, we found that relaxing this cutoff to 0.78 for the Sec proteome led to the inclusion of additional ESX-1 substrates EsxA and EspB, as well as the putative chaperone EspL. While we do not include these in our reported proteomes, we speculate that additional replicates—as well as use of various *Mtb* strains cultured under other conditions, as noted above— could strengthen these identifications. Overall, how EsxAB, EsxGH and other substrates and components of the ESX secretion system interact during cell wall transport remains to be further explored. Nevertheless, we note that recent data on ESX-1 suggest a hierarchical model in which, rather than being shuttled or channeled through the cell wall, substrates such as EsxB support their own export out of the cell^55, 64^ and our data lend additional credence to this model.

In summary protein tagging by APEX2 offers unprecedented accuracy in localizing proteins to subcellular compartments of *M. tuberculosis*, including the cell wall. Our identification of the cell wall proteome with this method is an important step towards understanding essential pathways within the mycobacterial cell envelope, including the transport of metabolites through the cell wall. APEX2 is a broadly useful tool for localizing individual proteins and proteomes in *Mtb* and future studies adapting this approach will likely expand the subcellular proteome inventories and enable the detailed exploration of cell wall processes.

## SIGNIFICANCE

All bacteria rely on specialized transport systems to mediate the import and export of essential metabolites through their cell envelopes. In mycobacteria, including the human pathogen *Mycobacterium tuberculosis* (*Mtb*), transport-associated proteins have been identified in the cytoplasm and plasma membrane, but how metabolites move through the cell wall is unknown. In the absence of sequence homology to proteins of known function, localization to the cell wall is arguably the strongest possible experimental evidence for cell wall function that can be achieved on a proteome-wide level. Here we validated APEX2 peroxidase-mediated protein tagging in live *Mtb* as a highly accurate method for localizing proteins to the cell wall or cytoplasm and identified the APEX2-tagged proteomes of these subcellular compartments. Our data show that substrates of the Type VII ESX protein secretion system are exposed to the periplasm during secretion. Type VII secretion is required for mycobacterial virulence, but the mechanism of protein export through the cell wall is entirely unknown. Our results lend support to a model in which secreted substrates support their own export into the extracellular matrix and suggest that ESX secretion proceeds by a mechanism distinct from that of other bacterial secretion systems.

## ACKNOWLEDGMENTS

We thank Mukshud Ahamed, Adrian Chen, Michael Li, Lauryn Martin, Uday Ganapathy, Jennifer Wang, Ian Winkeler for their contributions to preliminary experiments and data analyses and the Seeliger Lab for helpful discussions. We also thank Dheeraj Ramchandani for writing the Python script to convert UniProt identifiers to Rv gene identifiers. This work was supported by R35 GM119437 (J.C.S.) and R21 AI126044 (J.C.S.).

## AUTHOR CONTRIBUTIONS

Conceptualization: N.J., J.C.S. Data curation: J.A., B.U. Formal analysis: N.J., J.C.S., J.A., M.A., B.U. Funding acquisition: J.C.S. Investigation: N.J., M.L.P. Methodology: N.J., J.C.S. Project administration: J.C.S. Supervision: N.J., J.C.S., B.U. Validation: N.J., J.A., B.U. Visualization: N.J., J.C.S. Writing – original draft: N.J., J.C.S. Writing – review & editing: N.J., M.L.P., J.A., M.A., B.U., J.C.S.

## STAR METHODS

### RESOURCE AVAILABILITY

#### Lead contact

Further information and requests for resources and reagents should be directed to and will be fulfilled by the lead contact, Jessica Seeliger.

#### Materials availability

Plasmids generated in this study have been deposited in Addgene: pRibo-APEX2m (Addgene #176842) and pRibo-Sec-APEX2m (Addgene #176844).

#### Data and code availability

- All proteomics data are accessible at https://massive.ucsd.edu under MassIVE ID: MSVXXXXXX.
- All original code has been deposited at GitHub (https://github.com/dheerajramchandani/gene-name-fetcher) and is publicly available.
- Any additional information required to reanalyze the data reported in this paper is available from the lead contact upon request.

### EXPERIMENTAL MODEL DETAILS

All experiments were performed with the attenuated *Mycobacterium tuberculosis* (*Mtb*) strain H37Rv mc^2^6020 (*ΔlysA ΔpanCD*; gift of Dr. William Jacobs)^65^. *Mtb* was cultured in Middlebrook 7H9 medium (BD) supplemented with 10% v/v oleic acid-albumin-dextrose-catalase (OADC) supplement (BD), 0.5% v/v glycerol, 0.2% w/v casamino acids (VWR J851), 80 mg/L L-lysine, 24 mg/L pantothenate, and 0.025% v/v Tyloxapol or on Middlebrook 7H10 agar (BD) supplemented with 10% v/v OADC, 0.5% v/v glycerol, 0.2% casamino acids, 80 mg/L L-lysine, 24 mg/L pantothenate. Mtb strains harboring APEX2-expressing plasmids were selected on medium containing 25 μg/mL kanamycin. Mtb was cultured at 37 °C and in liquid medium with shaking at 110 rpm.

### METHOD DETAILS

#### Molecular cloning and transformation

The APEX2 sequence was optimized for codon usage in *Mtb* H37Rv using JCat (www.jcat.de)^66^. The resulting optimized sequence (APEX2m) was synthesized and cloned into the pUC57 plasmid (GeneWiz/Azenta) to yield pUC57-APEX2m. The APEX2m gene insert was amplified from pUC57-APEX2m using primers omlp492 (GCAACAAGATG<u>CATATG</u>GGCAAGAGCTACCCGACC) and olmp493 (GTTTTTGTTC<u>AAGCTT</u>GGCGTCGGCGAAGC) (restriction sites underlined) and the plasmid pRibo-BsaHind (Addgene #36251) was digested with BsaI and HindIII. Insert and digested vector were gel purified and assembled by In-Fusion cloning (Takara Bio) to yield the plasmid pRibo-APEX2m (Addgene #176842). The plasmids pRibo-Sec-APEX2 (Addgene #111699) and pUC57-APEX2m were digested with BamHI and ClaI. The vector backbone and APEX2m insert, respectively, were gel purified and ligated to yield pRibo-Sec-APEX2m (Addgene #176844). All constructs were confirmed by Sanger sequencing. The plasmids pRibo-APEX2m and pRibo-Sec-APEX2m were electroporated into *Mtb* and selected on medium containing kanamycin to yield the strains *Mtb*/APEX2m and *Mtb*/Sec-APEX2m.

#### APEX2-mediated labeling and enrichment of biotinylated proteins

Growth, APEX2-mediated labeling, and enrichment of biotinylated proteins was performed as reported^23^. Briefly, *Mtb*/APEX2m and *Mtb*/Sec-APEX2m were cultured in liquid medium to an optical density at 600 nm (OD_600_) of 0.8-1. *Mtb* was then subclutured to OD_600_ 0.25 in 50 mL (*Mtb*/APEX2m) or 250 mL (*Mtb*/Sec-APEX2m) with or without 2 mM theophylline and cultured for a further 72 h. Cells were harvested by centrifugation at 2500 x*g* for 10 min, resuspended at 10X concentration in 5 mL (*Mtb*/APEX2m) or 25 mL (*Mtb*/Sec-APEX2m) fresh medium containing 1 mM biotin-phenol (Iris Biotech), and incubated at 37 °C for 30 min. After harvesting by centrifugation, cells were resuspended in PBS and lysed by bead beating. Clarified lysates were then extracted with 1 % w/v dodecyl-maltoside at 4 °C for 1 h. Pre-washed neutravidin beads (Pierce PI29201) in PBS were added to lysates, incubated with gentle mixing at 22 °C for 1 h, and then washed with PBS. Enrichment and yield were confirmed by analyzing a fraction of the beads by SDS-PAGE followed by streptavidin Western blot (Streptavidin IR-Dye 680 LT; LI-COR Odyssey detection) and silver stain (Pierce), respectively.

#### Succinimidyl ester labeling of Mtb

*Mtb* was cultured in liquid medium to OD_600_ 0.8-1 and then subclutured to OD_600_ 0.25 in 250 mL further 72 h. Cells were harvested by centrifugation at 2500 x*g* for 10 min, washed twice with PBS with 0.5% Tween 80, and resuspended in 1/10^th^ the original volume (25 ml) PBS with 1 mM sulfo-NHS-biotin (Sulfo) or NHS-PEG-biotin (PEG) (Molecular Probes). Following a 30 min incubation at 4 °C, bacteria were washed three times with wash buffer (50 mM Tris-HCl, pH 8.0, 100 mM glycine, 0.01% Tween 80). After harvesting treated and washed cells by centrifugation, lysates were obtained and enriched for biotinylated proteins as above.

#### Western blot analysis of representative cell wall proteins

Total lysates and avidin-enriched fractions obtained above were normalized by protein concentration (BCA protein quantification kit; Pierce) or percent of the total beads, respectively. Samples were analyzed by SDS-PAGE and transferred to nitrocellulose. After blocking in PBS with 5% w/v bovine serum albumin and 0.1% Tween-20, membranes were probed with anti-LprG (supernatant from anti-Rv1411c clone B hybridoma, gift of Karen Dobos; equivalent antibody available also as NR-51133 from BEI Resources) or anti-Ag85 (BEI NR-13800, 1:5000 dilution) antibodies with gentle mixing at 4°C for 16 h. After washing three times with PBS with 0.1% Tween-20, membranes were incubated with anti-mouse IR Dye 280 LT or anti-rabbit IR Dye 800 LT secondary antibodies, respectively, (1:10000 dilution), for 1 h at 22 °C. After repeating the washes, the membranes were scanned (Odyssey Clx; LI-COR).

#### Identification of enriched proteins by label-free proteomics

On-bead digestion and peptide extraction were performed as previously described^67^. Briefly, proteins were reduced with dithiothreitol at 57 °C for 1 h (200 mM) and alkylated with iodoacetamide at 22 °C in the dark for 45 min (500 mM). Sequencing grade modified trypsin (Promega) was added to the sample for overnight digestion on a shaker at 22 °C (500 ng). The supernatant containing the peptides was removed, added to 200 µl R2 50 µM Poros Beads and shaken for 3 h at 4 °C. The beads were loaded onto equilibrated C18 ziptips (Millipore) using a microcentrifuge, the sample rinsed three times with 0.1% TFA, and further washed with 0.5% acetic acid. Peptides were eluted with 40% acetonitrile in 0.5% acetic acid followed by 80% acetonitrile in 0.5% acetic acid. The organic solvent was removed using a concentrator (SpeedVac) and the desalted peptides reconstituted in 0.5% acetic acid and stored at −80°C until analysis.

One-tenth of the total sample was analyzed by LC-MS using an EASY-nLC 1200 (Thermo Scientific) in-line with a Q Exactive mass spectrometer (Thermo Fisher Scientific) using a 1 h gradient (Solvent A: 2% acetonitrile, 0.5% acetic acid; Solvent B: 80% acetonitrile, 0.5% acetic acid). High resolution full MS spectra were acquired with a resolution of 70,000, an AGC target of 1 x 10^6^, a maximum ion time of 120 ms, and scan range of 400 to 1500 *m/z*. Following each full MS, twenty data-dependent high resolution HCD MS/MS spectra were acquired. All MS/MS spectra were collected using the following instrument parameters: resolution of 17,500, AGC target of 5 x 10^4^, maximum ion time of 120 ms, one microscan, 2 *m/z* isolation window, fixed first mass of 150 *m/z*, and NCE of 27. MS/MS spectra were searched against a UniProt *Mycobacterium tuberculosis* database using Sequest within Proteome Discoverer 1.4. The results were filtered using a 1% false discovery rate peptide and protein cut off searched against a decoy database using Percolator (http://percolator.ms/)^68^ and proteins had to have at least 2 unique peptides to be considered. A SAINT Express analysis^69^ was performed using the samples without theophylline as control to rank the identified proteins based on their likelihood to be bona fide enriched (i.e., biotinylated) in a theophylline-dependent (and thus APEX2-dependent) manner (**Table S1**). Hits were defined as proteins with SAINT scores >0.9. Since proteins of the WXG100 family (including Esx substrates) have high sequence redundancy, additional analysis confirmed the unique identification of individual substrates (EsxA-EsxL, EsxN-EsxW). Among Esx substrates identified in any APEX2 proteome, only EsxL/EsxN could not be uniquely assigned. UniProt identifiers in the resulting proteomic datasets were used to obtain the corresponding Rv reference numbers using a custom Python script (https://github.com/dheerajramchandani/gene-name-fetcher).

#### Prediction of protein export signals and structures

Using the UniProt identifiers, FASTA sequence files for the 490 proteins detected from *Mtb* expressing APEX2 and 254 proteins detected from *Mtb* expressing Sec-APEX2 were obtained from the UniProt database (http://www.uniprot.org). These FASTA files were then used as input to identify predicted signal sequences using SignalP (version 6.0)^70^ and predicted transmembrane helices using DeepTMHMM (version 1.0.12)^71^. The presence of a consensus ESX secretion signal as reported by Daleke et al^72^ was detected using ScanProSite^73^. The Alphafold plugin within the UniProt database was used to obtain predicted structures for selected proteins and to generate customized protein structure views with tyrosines highlighted.

### QUANTIFICATION

Densitometry analysis to quantify enrichment of LprG and Ag85 was performed with ImageJ^74^. Signal intensities from output samples (post enrichment) were normalized to input samples. The normalized values were then used to calculate the ratio of test (with APEX2 expression) to control (without APEX2 expression).

## SUPPLEMENTAL INFORMATION

**Supplemental Figure S1.** Biotinylation of proteins in Mtb by Cyt-APEX2 or Sec-APEX2 (additional replicates)

**Supplemental Figure S2.** Validation of compartment-specific labeling by Cyt-APEX2 or Sec-APEX2 (additional replicate)

**Supplemental Figure S3.** APEX2-mediated labeling predicts topology of transmembrane proteins

**Supplemental Table S1.** List of proteins and associated peptides enriched from *Mtb* lysates **Supplemental Table S2.** Comparison of proteomes detected by APEX2 (Cyt), Sec-APEX2 (Sec), or NHS-PEG-biotin (NHS) labeling to reported cell wall proteomes

**Supplemental Table S3.** Analysis of the NHS-labeled (NHS) proteome for proteins with predicted cytosolic localization and/or function

**Supplemental Table S4.** Analysis of the Sec-APEX2 (Sec) proteome for proteins with predicted cytosolic localization and/or function

**Supplemental Table S5.** Comparison of proteins reported in cell wall proteomes that have known or predicted cytosolic functions with the APEX2 (Cyt) and Sec-APEX2 (Sec) proteomes

**Supplemental Table S6.** Comparison of the EXIT proteome to APEX2 (Cyt) and Sec-APEX2 (Sec) proteomes

**Supplemental Table S7.** Comparison of proteins reported in Kavunja et al. with the APEX2 (Cyt) and Sec-APEX2 (Sec) proteomes

**Supplemental Table S8.** Identification of divisome/elongasome proteins in the Cyt-APEX2 (Cyt) and Sec-APEX2 (Sec) proteomes

**Supplemental Table S9.** Comparison of culture filtrate (CF) proteomes with the Cyt-APEX2 (Cyt) and Sec-APEX2 (Sec) proteomes

