## Supplemental Figures S1-S3 for "Cell wall proteomics in live *Mycobacterium tuberculosis* uncovers exposure of ESX substrates to the periplasm"

**SUPPLEMENTAL INFORMATION**

**Supplemental Figure S1.** Biotinylation of proteins by Cyt-APEX2 or Sec-APEX2 (additional replicates)

**Supplemental Figure S2.** Validation of compartment-specific labeling by Cyt-APEX2 or Sec-APEX2 (additional replicate)

**Supplemental Figure S3.** APEX2-mediated labeling predicts topology of transmembrane proteins

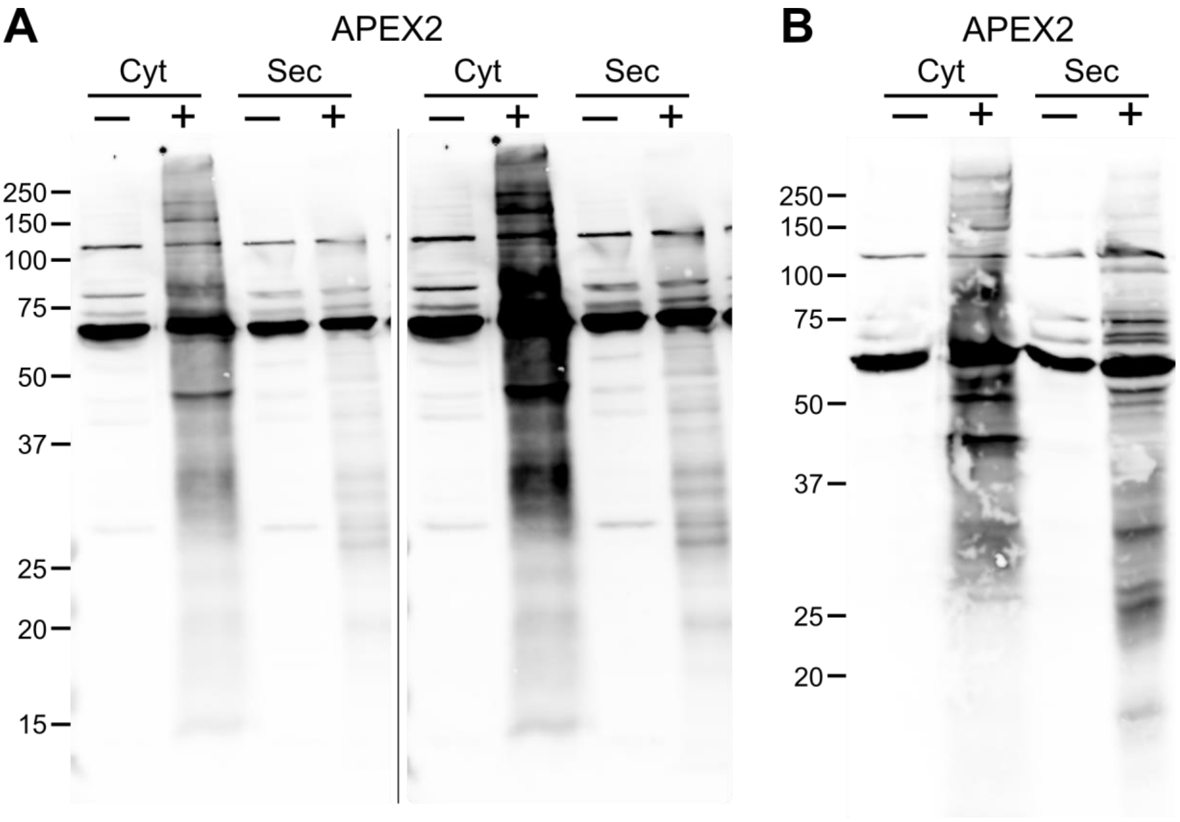


**Supplemental Figure S1. Biotinylation of proteins by Cyt-APEX2 or Sec-APEX2 (additional replicates).** *Mtb* encoding inducible Cyt-APEX2 or Sec-APEX2 was cultured without (–) or with (+) theophylline and subjected to the labeling protocol with biotin-phenol. (A) and (B) show 2 additional replicates of Figure 1A. In (A) the image on the right is the same blot contrasted to highlight Sec-APEX2-dependent biotinylation. For (B) contrasting was not necessary to visualize labeling by Sec-APEX2 clearly. For both blots, biotinylation in crude lysates were detected with streptavidin.

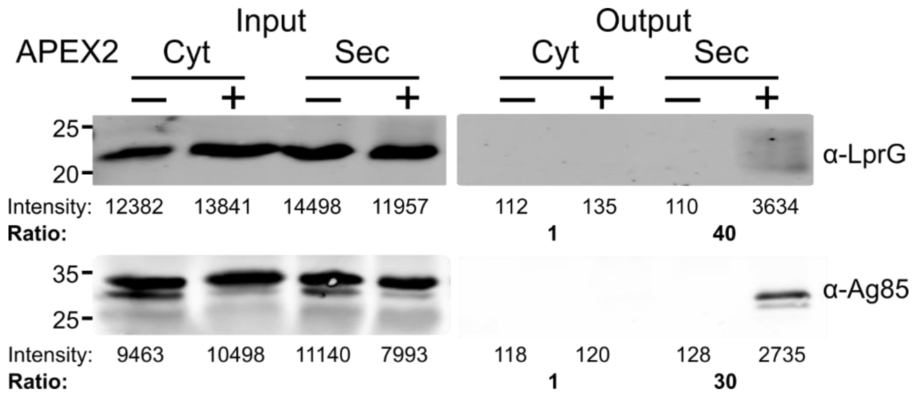


**Supplemental Figure S2. Validation of compartment-specific labeling by Cyt-APEX2 or Sec-APEX2 (additional replicate).** Biotinylated proteins were enriched from crude lysates obtained from *Mtb* encoding inducible Cyt-APEX2 or Sec-APEX2 and cultured without (─) or with (+) theophylline before being subjected to the labeling protocol with biotin-phenol. The fold increase in biotinylation upon induction of APEX2 or treatment with NHS (“Ratio”) was calculated by taking the ratio of the + / − theophylline output intensities after normalizing to the corresponding inputs.

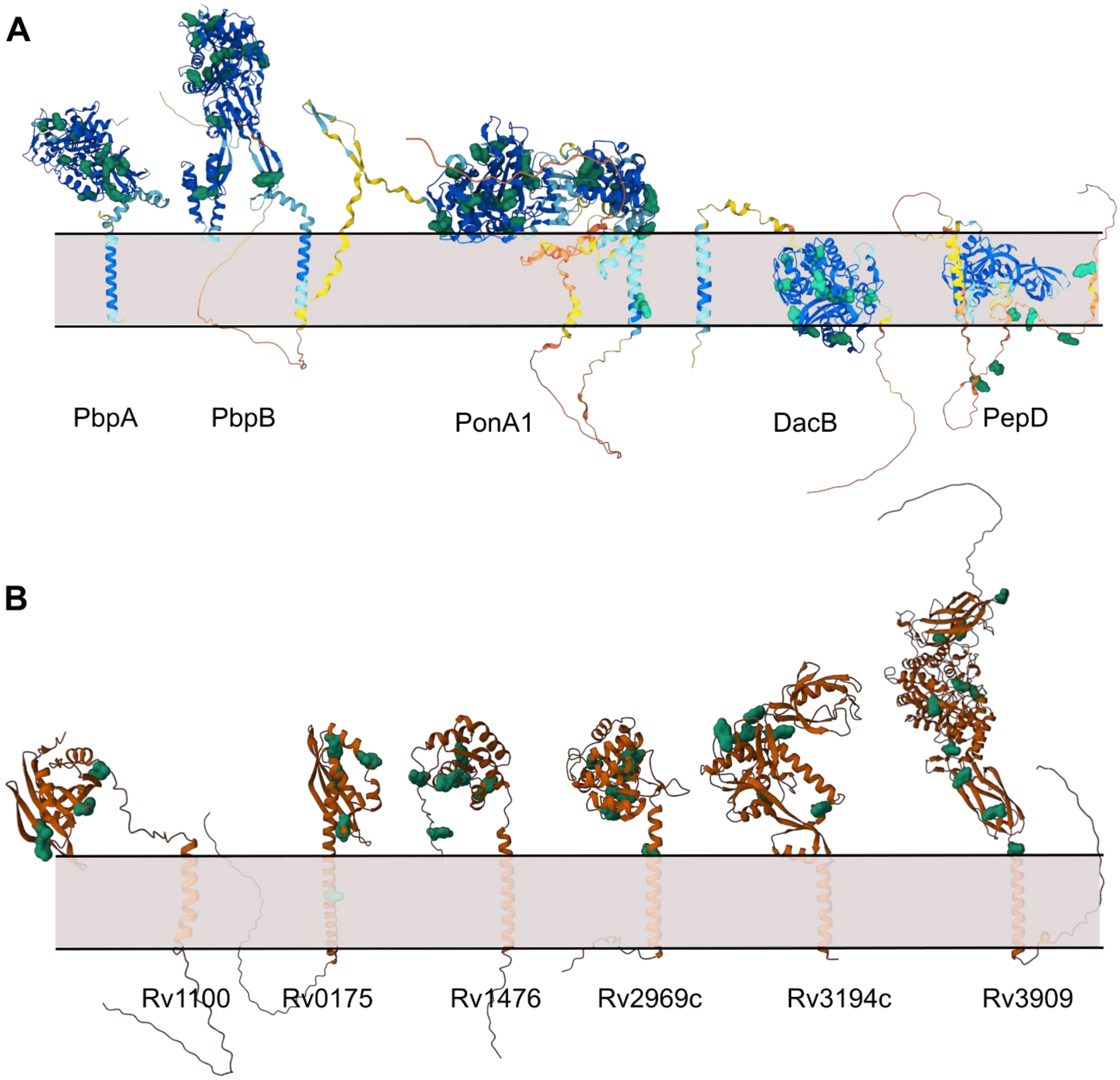


**Supplemental Figure S3. APEX2-mediated labeling predicts topology of transmembrane proteins.** (A) Predicted structures of PbpA, PbpB, PonA1, DacB, and PepD (HtrA2) were obtained from the AlphaFold plugin within Uniprot (uniprot.org). Tyrosine residues are highlighted in green. (B) Predicted structures of uncharacterised proteins were downloaded from AlphaFold (https://alphafold.ebi.ac.uk/) and tyrosines highlighted as above using the Mol* 3D Viewer plugin within the PDB website(https://www.rcsb.org/3d-view). Structures are shown oriented relative to the predicted transmembrane helix and based on tyrosine location and detection in the Cyt or Sec proteome.
